# No evidence that inbreeding avoidance is up-regulated during the ovulatory phase of the menstrual cycle

**DOI:** 10.1101/192054

**Authors:** Iris J. Holzleitner, Julie C. Driebe, Ruben C. Arslan, Amanda C. Hahn, Anthony J. Lee, Kieran J. O’Shea, Tanja M. Gerlach, Lars Penke, Benedict C. Jones, Lisa M. DeBruine

## Abstract

Mate preferences and mating-related behaviors are hypothesized to change over the menstrual cycle in ways that function to increase reproductive fitness. Results of recent large-scale studies suggest that many of these hormone-linked behavioral changes are less robust than was previously thought. One specific hypothesis that has not yet been subject to a large-scale test is the proposal that women’s preference for associating with male kin is down-regulated during the ovulatory (high-fertility) phase of the menstrual cycle. Consequently, we used a longitudinal design to investigate the relationship between changes in women’s steroid hormone levels and their perceptions of faces experimentally manipulated to possess kinship cues (Study 1). Analyses suggested that women viewed men’s faces displaying kinship cues more positively (i.e., more attractive and trustworthy) when estradiol-to-progesterone ratio was high. Since estradiol-to-progesterone ratio is positively associated with conception risk during the menstrual cycle, these results directly contradict the hypothesis that women’s preference for associating with male kin is down-regulated during the ovulatory (high-fertility) phase of the menstrual cycle. Study 2 employed a daily diary approach and found no evidence that women reported spending less time in the company of male kin or thought about male kin less often during the fertile phase of the menstrual cycle. Thus, neither study found evidence that inbreeding avoidance is up-regulated during the ovulatory phase of the menstrual cycle.

---

Many researchers have proposed that during the ovulatory (i.e., high-fertility) phase of the menstrual cycle, women’s preferences for potential mates who will increase their reproductive fitness will strengthen and/or that women’s aversions to potential mates who will decrease their reproductive fitness will strengthen (see Gildersleeve et al., 2014; Gangestad & Thornhill 2008; Jones et al., 2008 for reviews). Increased attraction to men displaying putative good-fitness cues (Gangestad et al., 2004; Gangestad et al., 2007; Penton-Voak et al., 1999; Penton-Voak & Perrett, 2000) during the ovulatory phase of the menstrual cycle are particularly high-profile (but not the only) examples of evidence that is widely cited for this claim.

Recently, however, the robustness of the evidence for ovulatory shifts in women’s mate preferences has been called into question. For example, two different meta-analyses of this literature drew very different conclusions about the robustness of the evidence for ovulatory shifts in women’s mate preferences (Gildersleeve et al., 2014; Wood et al., 2004). Researchers have also highlighted several potentially important methodological limitations of studies on this topic (Blake et al., 2016; Gangestad et al., 2016; Jones et al., 2018a).

First, many researchers have emphasized that the majority of studies reporting significant ovulatory shifts in these behaviors are badly underpowered (Gangestad et al., 2016; Jones et al., 2018a). In combination with publication bias, this issue means that many of the published effects are likely to be false positives.

Second, many studies in this literature have employed between-subjects (i.e., cross-sectional) designs, which are ill-suited for testing subtle ovulatory shifts in behaviors that have substantial between-subject variance (Gangestad et al., 2016; Jones et al., 2018a). Importantly, large-scale within-subject (i.e., longitudinal) studies that used more objective methods to assess women’s hormonal status (e.g., measuring sex hormones from saliva) have generally not replicated previously reported findings for ovulatory shifts in mate preferences (Jones et al., 2018a; Jünger et al., 2018a; Jünger et al., 2018b; Marcinkowska et al., 2018).

Third, studies have typically used self-report methods to assess position in the menstrual cycle (e.g., self-reported number of days since last period of menstrual bleeding at time of testing). Empirical studies suggest these are imprecise and prone to bias (Blake et al., 2016), although this may not be a problem in longitudinal studies with very large samples (e.g., Arslan et al., 2018).

Behaviors aimed at reducing opportunities for inbreeding to occur are predicted to increase around ovulation (Lieberman et al., 2011) but have yet to be the focus of large-scale, rigorous tests. To date, the best evidence for ovulatory shifts in inbreeding-avoidance comes from Lieberman et al. (2011). In a longitudinal study of 48 women’s mobile phone records from one menstrual cycle, Lieberman et al. reported that women called their fathers less frequently, and spoke to them for less time when they did call them, during the high-fertility phase of the menstrual cycle than when fertility was low. Because Lieberman et al. observed no such change in women’s frequency or duration of calls to their mothers, they interpreted these results as evidence for adaptations that function to reduce opportunities for inbreeding to occur around ovulation. Consistent with Lieberman et al.’s findings, DeBruine et al. (2005) found that women showed stronger preferences for faces manipulated to possess kinship cues during the luteal (low-fertility) phase of the menstrual cycle than during the ovulatory phase in a cross-sectional study of 71 women. However DeBruine et al. (2005) also found that preferences for cues of kinship in women’s, but not men’s, faces were related to women’s progesterone level, but not estimated fertility. Both progesterone and fertility were estimated by converting reported menstrual cycle day to progesterone and conception risk values using actuarial tables.

Researchers have recently emphasized the importance of replicating cyclic shifts in behaviors that have not yet been the target of large-scale replications, including inbreeding avoidance (Jones et al., in press). Thus, we revisited the claim of hormonal regulation of inbreeding-avoidance behaviors.

In Study 1, we examined this claim in a large-scale longitudinal study of the relationship between women’s (N=199) salivary hormone levels and their responses to kinship cues in faces. Following previous studies of responses to facial kinship cues (DeBruine, 2002, 2004, 2005; DeBruine et al., 2005), we experimentally manipulated male and female face images to be more or less similar in shape to our participants and assessed the effects of this manipulation on perceptions of attractiveness and trustworthiness. Previous research has shown that this image manipulation can reliably tap inbreeding-avoidance behaviors. For example, women show aversions to opposite-sex faces with similar shape characteristics to their own when assessing men for exclusively sexual relationships, such as one-night stands, but not when assessing their trustworthiness (DeBruine, 2005). Moreover, such effects are not due to feminization of opposite-sex faces when increasing self-resemblance to female participants (DeBruine, 2005). Further evidence that people respond to this image manipulation in ways consistent with it functioning as a kinship cue comes from studies showing that people are more likely to cooperate with people with similar face-shape characteristics (DeBruine, 2002) and perceive them to be more trustworthy (DeBruine, 2005).

The ovulatory phase of the menstrual cycle is characterized by the combination of high estradiol and low progesterone (Gangestad & Haselton, 2015). Thus, if Lieberman et al. (2011) are correct that ovulation increases inbreeding-avoidance behaviors, we would expect preferences for self-resembling male, but not self-resembling female, faces to decrease when estradiol is high and progesterone is simultaneously low.

In Study 2, we tested for hormonal regulation of inbreeding-avoidance behaviors by investigating whether women’s reported frequency of contact with and frequency of thought about male kin increased when conception risk was high.

## Study 1

Study 1 investigated whether women’s responses to kinship cues in faces track changes in estradiol and progesterone.

## Methods

### Participants

We tested 205 heterosexual women (mean age=21.5 years, SD=3.3 years) who reported that they were not using any form of hormonal contraceptive (i.e., reported having natural menstrual cycles). Participants completed up to three blocks of test sessions. Each of the three blocks of test sessions consisted of five weekly test sessions. Women participated as part of a large study of possible effects of steroid hormones on women’s behavior (Jones et al., 2018a, 2018b, 2018c). The data analysed here are all responses from blocks of test sessions where women were not using any form of hormonal contraceptive and completed the face-judgment task in at least one test session. One hundred and seventy-two women had completed four or more test sessions and 41 of these women completed nine test sessions. Thirty-three women completed fewer than five test sessions.

### Procedure

In the first test session, a full-face photograph of each woman was taken under standardized photographic conditions. Camera-to-head distance was held constant. These photographs were used to manufacture self-resembling faces using the same methods as previous research (DeBruine, 2002, 2004, 2005; DeBruine et al., 2005). Self-resembling faces were created by applying 50% of the shape difference between each participant’s face and a same-sex (i.e. female) prototype face to same-sex and opposite-sex prototypes, to produce same-sex and opposite-sex self-resembling faces. Importantly, this method for manipulating self-resemblance in opposite-sex faces (DeBruine, 2004) avoids the feminization of male stimulus faces that occurs when simply blending self and opposite-sex faces. Male and female comparison stimuli that resembled none of the participants were manufactured in the same way using images of ten women who did not participate in the study. As in previous research on responses to self-resembling faces (DeBruine, 2002, 2004), image manipulations were carried out using specialist computer graphic software (DeBruine, 2018; Tiddeman et al., 2001). Example stimuli are shown in Figure 1.

**Figure 1.**
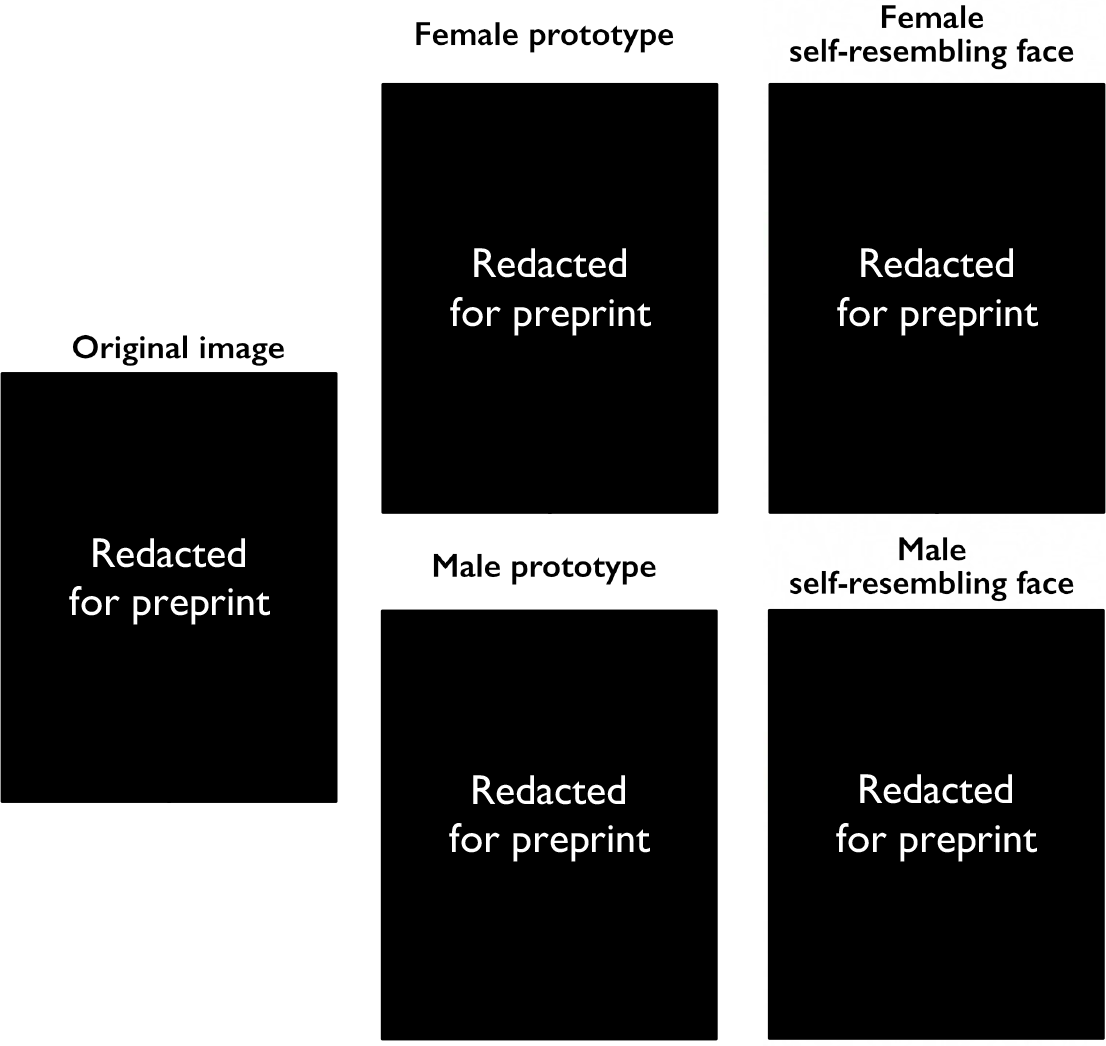
Self-resembling stimulus faces were created by applying 50% of the difference in shape between an individual’s face and the female prototype to both female and male prototype faces.

In all subsequent test sessions (all test sessions after the first), each woman completed a face-judgment task in which they were presented 20 pairs of faces. Ten of these pairs consisted of a self-resembling face and a comparison face. The other ten pairs consisted of a non-resembling face (constructed from another randomly selected age-matched woman participating in the study) and the same comparison faces. This method allows us to compare judgments of self-resembling faces to judgments of non-resembling faces, while keeping equal the number of times self- and non-resembling faces are presented.

Participants were instructed to click on the face in each pair that they thought looked more attractive or, in a separate block of trials, more trustworthy. Trial order and the side of the screen on which any given image was presented were fully randomized. In each test session, each woman completed the face-judgment task four times. In the first version, they were presented female faces and judged attractiveness.

In the second version, they were presented female faces and judged trustworthiness. In the third version, they were presented male faces and judged attractiveness. In the fourth version, they were presented male faces and judged trustworthiness. The order in which participants completed these versions of the face-judgment task was fully randomized.

For each version of the face-judgment task, we calculated a self-resemblance bias score by subtracting the number of times the non-resembling faces were chosen (out of 10) from the number of times the self-resembling faces were chosen (out of 10). These four scores were calculated separately for each participant in each test session. Positive scores indicated a bias towards self-resembling (versus non-resembling) faces and higher scores indicated self-resembling faces were perceived as more attractive or trustworthy than control faces.

### Saliva samples

Participants also provided a saliva sample via passive drool (Papacosta & Nassis, 2011) in each test session. Participants were instructed to avoid consuming alcohol and coffee in the 12 hours prior to participation and avoid eating, smoking, drinking, chewing gum, or brushing their teeth in the 60 minutes prior to participation. Each woman’s test sessions took place at approximately the same time of day to minimize effects of diurnal changes in hormone levels (Veldhuis et al., 1988; Bao et al., 2003).

Saliva samples were frozen immediately and stored at −32°C until being shipped, on dry ice, to the Salimetrics Lab (Suffolk, UK) for analysis, where they were assayed using the Salivary 17β-Estradiol Enzyme Immunoassay Kit 1-3702 (M=2.82 pg/mL, SD=1.03 pg/mL, sensitivity=0.1 pg/mL, intra-assay CV=7.13%, inter-assay CV=7.45%) and Salivary Progesterone Enzyme Immunoassay Kit 1-1502 (M=157.2 pg/mL, SD=104.9 pg/mL, sensitivity=5 pg/mL, intra-assay CV=6.20%, inter-assay CV=7.55%). Hormone levels more than three standard deviations from the sample mean for that hormone or where Salimetrics indicated levels were outside the sensitivity range of their relevant ELISA were excluded from the dataset (~0.1% of hormone measures were excluded for these reasons). The descriptive statistics given above do not include these excluded values and do not include statistics for the first test session where women did not complete the face-judgment task. Values for each hormone were centered on their subject-specific means to isolate effects of within-subject changes in hormones and were scaled so the majority of the distribution for each hormone varied from −.5 to .5. This was done simply to facilitate calculations in the linear mixed models. Since hormone levels were centered on their subject-specific means, women with only one value for a hormone could not be included in these analyses.

### Analyses and results

Linear mixed models were used to test for possible effects of hormonal status on responses on the face-judgment task. Analyses were conducted using R version 3.5.1 (R Core Team, 2018), with lme4 version 1.1-18-1 (Bates et al., 2014) and lmerTest version 3.0-1 (Kuznetsova et al., 2017). Random slopes were specified maximally following Barr et al. (2013) and Barr (2013). Data files and analysis scripts are publicly available at https://osf.io/wnhma/.

The models we used to investigate hormonal regulation of responses to kinship cues in faces are identical to those we have used previously to test for hormonal regulation of women’s masculinity preferences (Jones et al., 2018a), disgust sensitivity (Jones et al., 2018b), and sexual desire (Jones et al., 2018c). Face sex was effect coded (−.5 = female, +.5 = male), as was judgment type (−.5 = attractiveness, +.5 = trustworthiness). Self-resemblance bias scores (−10 to +10) were the dependent variable. Note that women with only a single test session where they completed the face-judgment task and had valid estradiol and progesterone levels cannot be included in these longitudinal analyses (n=6). Thus, data from 199 women were included in these analyses.

The first model (Model 1) we tested included estradiol (scaled and centered), progesterone (scaled and centered), estradiol-to-progesterone ratio (scaled and centered), face sex, and judgment type as predictors, as well as all possible two-way and three-way interactions among these predictors. Full results for this analysis are shown in Table 1. The intercept was positive and significant (estimate=0.43, 95% CI = [0.15, 0.72], p=.003), indicating that women chose self-resembling faces more often than would be predicted by chance. There was a significant positive effect of progesterone (estimate=0.63, 95% CI = [0.13, 1.13], p=.015) and a significant negative effect of estradiol (estimate=−0.76, 95% CI = [−1.31, −0.2], p=.008), indicating that self-resemblance-bias scores tracked changes in both progesterone and estradiol. Although self-resemblance bias scores tended to be higher for judgments of female faces than male faces, this effect of face sex was not significant (estimate=−0.21, 95% CI = [−0.43, 0.01], p=.068). The main effect of estradiol-to-progesterone ratio (estimate=0.34, 95% CI = [−0.11, 0.78], p=.159) was not significant, but the interaction between face sex and estradiol-to-progesterone ratio was significant (estimate=0.71, 95% CI = [0.02, 1.4], p=.044). However, self-resemblance bias scores for male faces were higher when estradiol-to-progesterone ratio was high (see Figure 2), which is in the opposite direction to the prediction that self-resemblance bias scores for male faces will decrease when conception risk is high. Self-resemblance bias scores for female faces did not appear to be related to estradiol-to-progesterone ratio (see Figure 2). No other effects were significant or close to being significant.

**Table 1.**
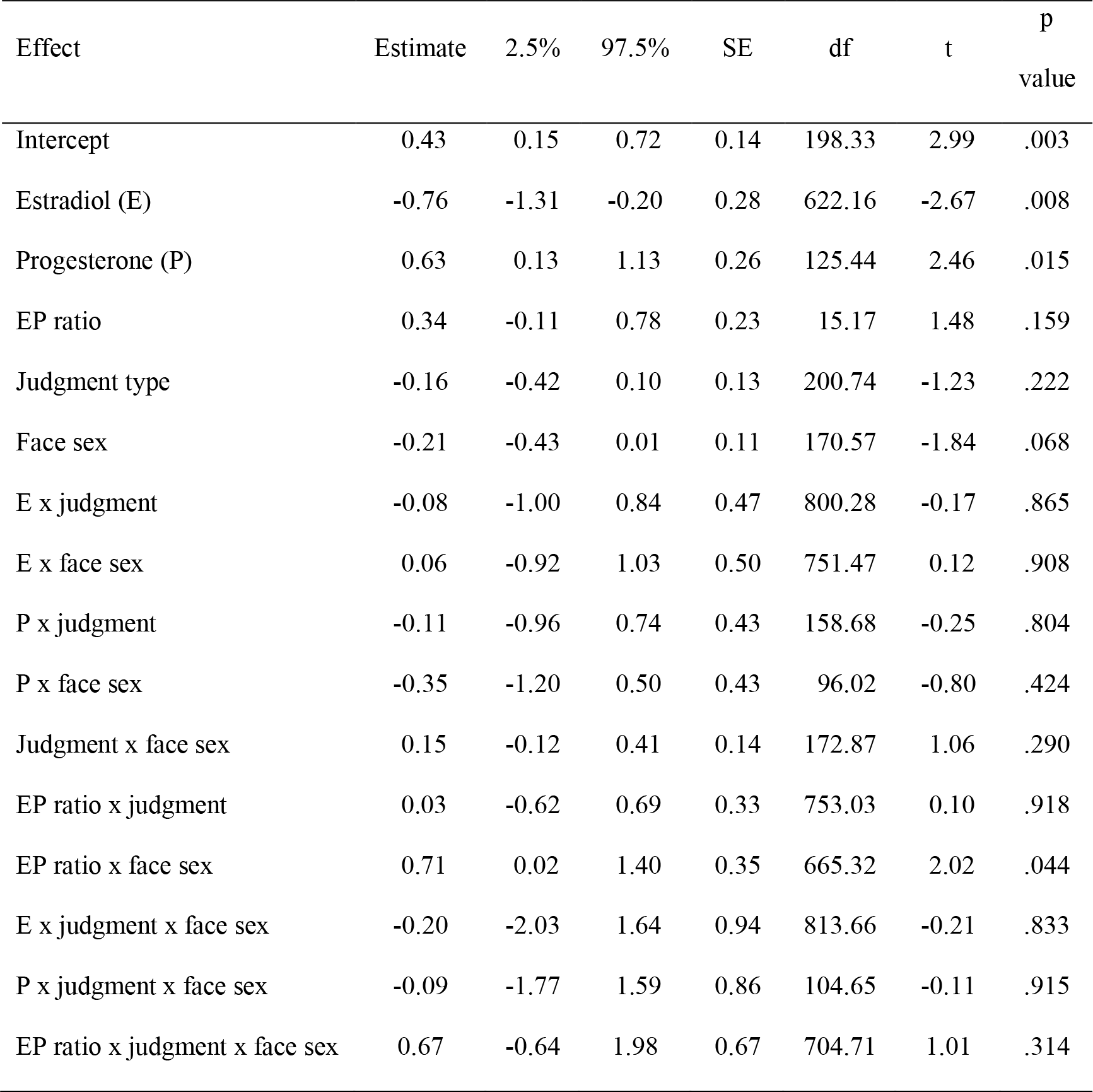
The effect of estradiol, progesterone and estradiol-to-progesterone ratio on self-resemblance bias scores (Model 1)

| Effect | Estimate | 2.5% | 97.5% | SE | df | t | p value |
| --- | --- | --- | --- | --- | --- | --- | --- |
| Intercept | 0.43 | 0.15 | 0.72 | 0.14 | 198.33 | 2.99 | .003 |
| Estradiol (E) | -0.76 | -1.31 | -0.20 | 0.28 | 622.16 | -2.67 | .008 |
| Progesterone (P) | 0.63 | 0.13 | 1.13 | 0.26 | 125.44 | 2.46 | .015 |
| EP ratio | 0.34 | -0.11 | 0.78 | 0.23 | 15.17 | 1.48 | .159 |
| Judgment type | -0.16 | -0.42 | 0.10 | 0.13 | 200.74 | -1.23 | .222 |
| Face sex | -0.21 | -0.43 | 0.01 | 0.11 | 170.57 | -1.84 | .068 |
| E x judgment | -0.08 | -1.00 | 0.84 | 0.47 | 800.28 | -0.17 | .865 |
| E x face sex | 0.06 | -0.92 | 1.03 | 0.50 | 751.47 | 0.12 | .908 |
| P x judgment | -0.11 | -0.96 | 0.74 | 0.43 | 158.68 | -0.25 | .804 |
| P x face sex | -0.35 | -1.20 | 0.50 | 0.43 | 96.02 | -0.80 | .424 |
| Judgment x face sex | 0.15 | -0.12 | 0.41 | 0.14 | 172.87 | 1.06 | .290 |
| EP ratio x judgment | 0.03 | -0.62 | 0.69 | 0.33 | 753.03 | 0.10 | .918 |
| EP ratio x face sex | 0.71 | 0.02 | 1.40 | 0.35 | 665.32 | 2.02 | .044 |
| E x judgment x face sex | -0.20 | -2.03 | 1.64 | 0.94 | 813.66 | -0.21 | .833 |
| P x judgment x face sex | -0.09 | -1.77 | 1.59 | 0.86 | 104.65 | -0.11 | .915 |
| EP ratio x judgment x face sex | 0.67 | -0.64 | 1.98 | 0.67 | 704.71 | 1.01 | .314 |

**Figure 2.**
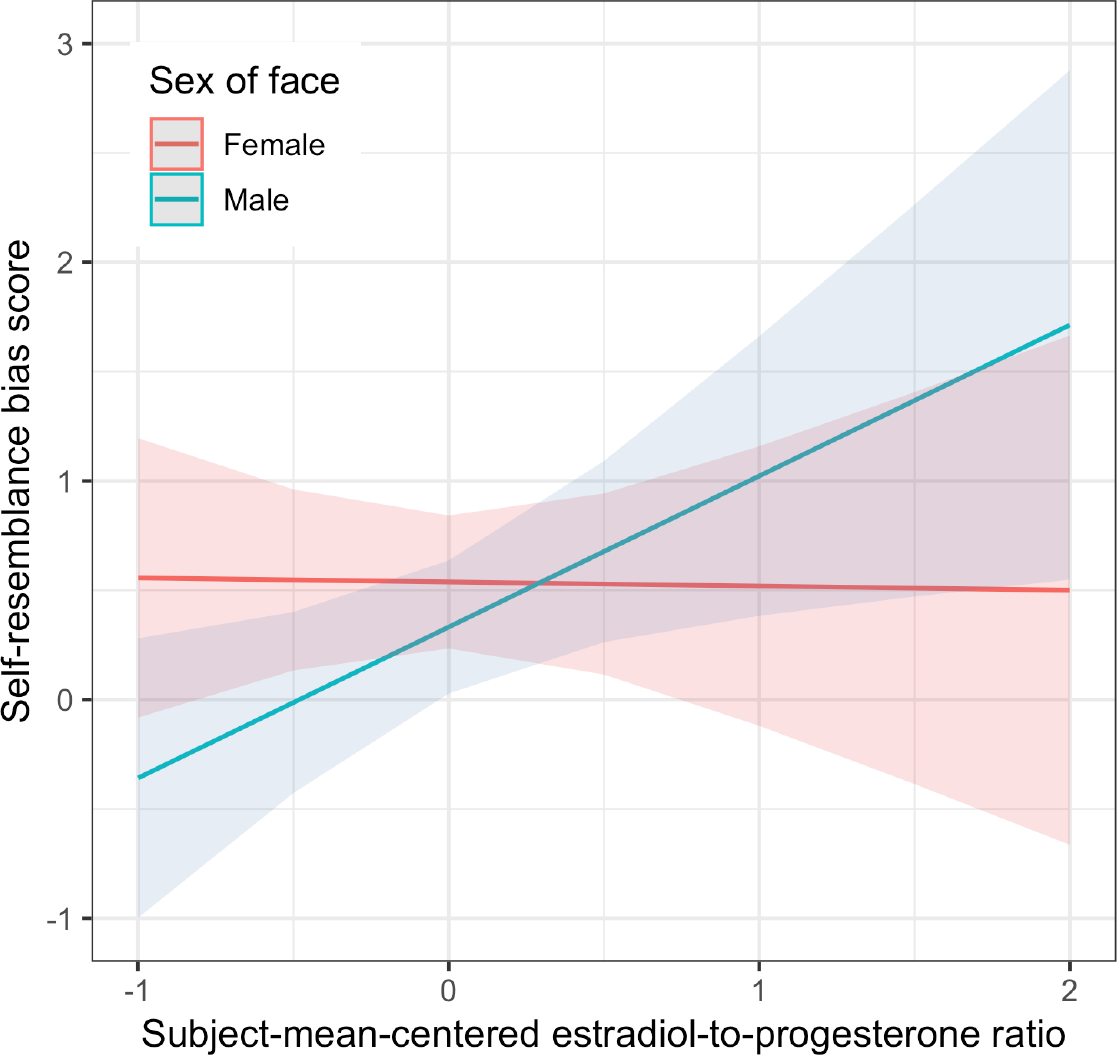
The effect of estradiol-to-progesterone ratio on self-resemblance bias scores. Self-resemblance bias scores for male faces were higher when estradiol-to-progesterone ratio was high, while self-resemblance bias scores for female faces appeared to be unrelated to estradiol-to-progesterone ratio.

The second model (Model 2) we tested included estradiol (scaled and centered), progesterone (scaled and centered), face sex, and judgment type as predictors, as well as all possible two-way, three-way, and four-way interactions among these predictors. This model produced convergence warnings with both the bobyqa or Nelder-Mead optimizers (see https://osf.io/wnhma/). To facilitate convergence, we ran a simplified model, in which only random slopes for the highest-order interaction and the interaction among face sex, estradiol, and progesterone were included. This model produced identical results when run with bobyqa and Nelder-Mead optimizers (see https://osf.io/wnhma/). Full results for this analysis are shown in Table 2. Replicating the results reported for Model 1, the intercept was significant (estimate=0.44, 95% CI = [0.15, 0.72], p=.003), there was a significant positive effect of progesterone (estimate=0.43, 95% CI = [0.03, 0.83], p=.033), and a significant negative effect of estradiol (estimate=−0.63, 95% CI = [−1.17, −0.08], p=.024). Self-resemblance bias scores were higher for judgments of female faces than male faces and this effect of face sex was significant in this analysis (estimate=−0.22, 95% CI = [−0.36, −0.08], p=.003). The effect of judgment type was significant in this analysis (estimate=−0.18, 95% CI = [−0.33, −0.04], p=.012), indicating that self-resemblance bias scores were greater for attractiveness judgments than trustworthiness judgments. In this analysis, the interaction between progesterone and face sex was significant (estimate=−0.84, 95% CI = [−1.63, −0.06], p=.035) and showed that the positive effect of progesterone on self-resemblance scores was greater for female than male faces (see Figure 3). No other effects were significant or close to being significant. Of note, neither the three-way interaction of progesterone, estradiol and sex of face (estimate=1.79, 95% CI = [−2.83, 6.42], p=.447), nor the two-way interaction of progesterone and estradiol (estimate=−0.77, 95% CI = [−3.43, 1.9], p=.573) were significant. The pattern of results observed for the simplified model are very similar to those observed in the full models that did not converge (see https://osf.io/wnhma/).

**Table 2.**
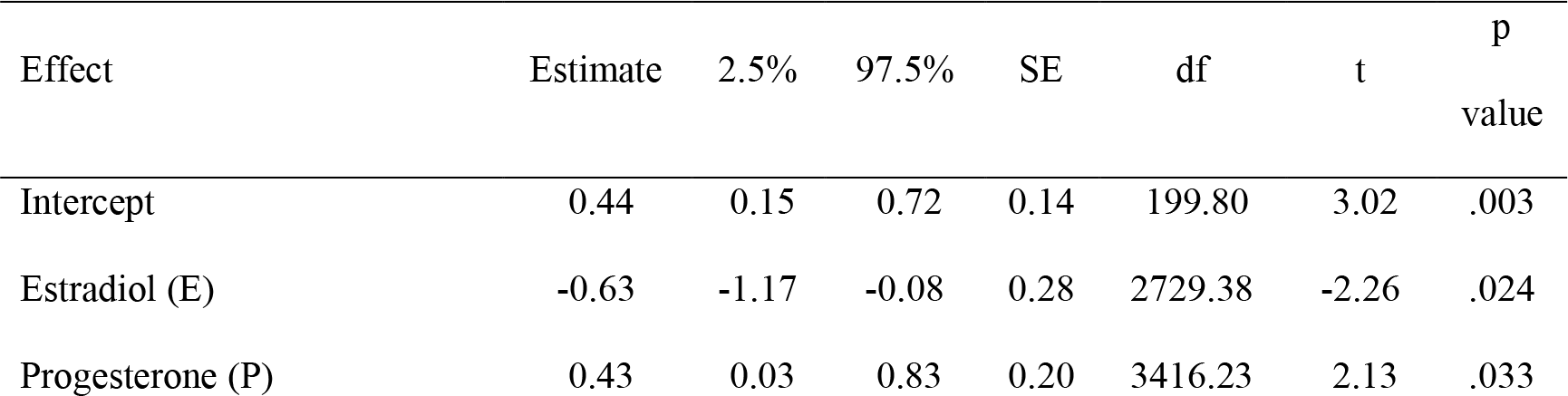
The effect of estradiol, progesterone and the interaction of estradiol and progesterone on self-resemblance bias scores (Model 2)

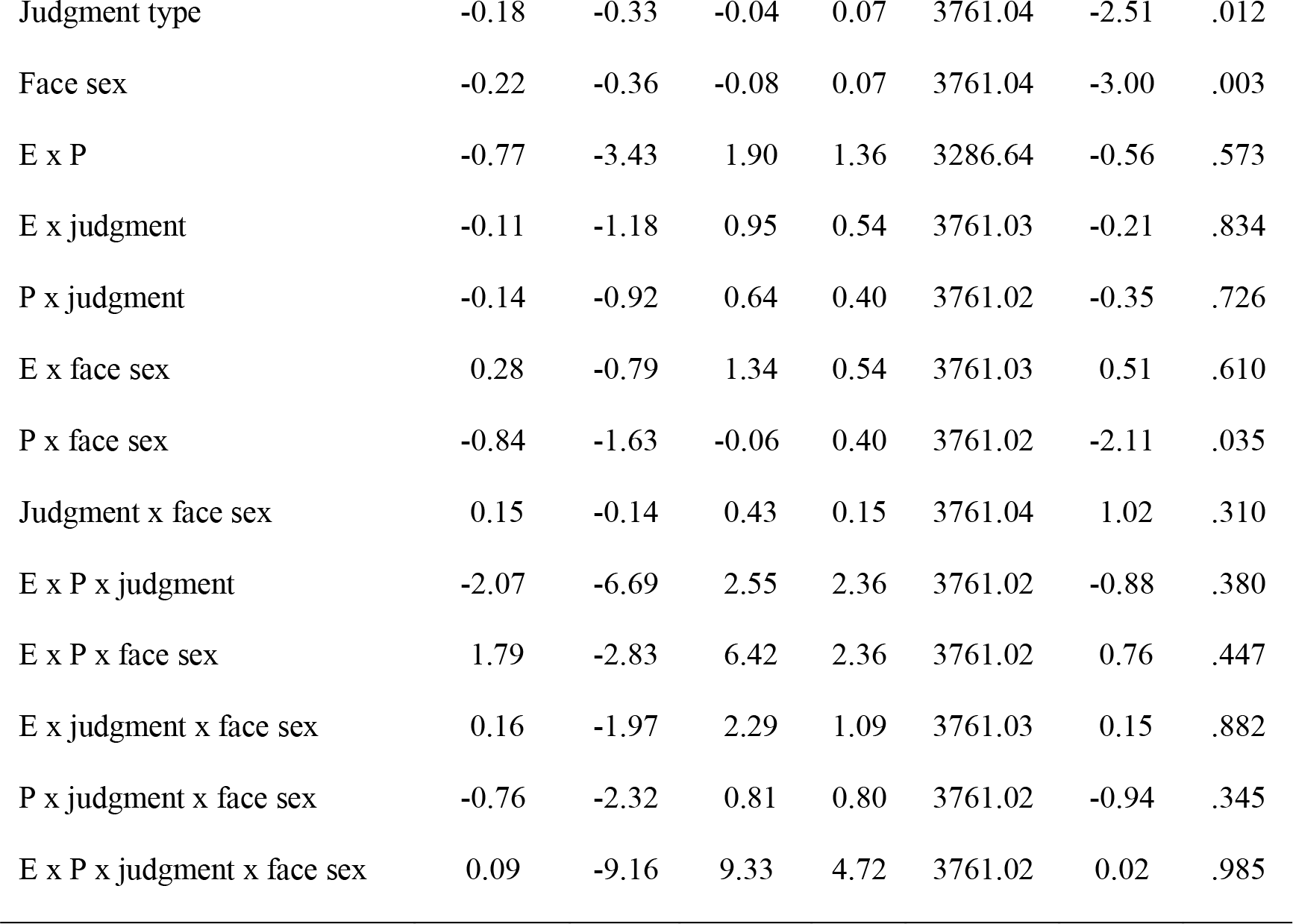

**Figure 3.**
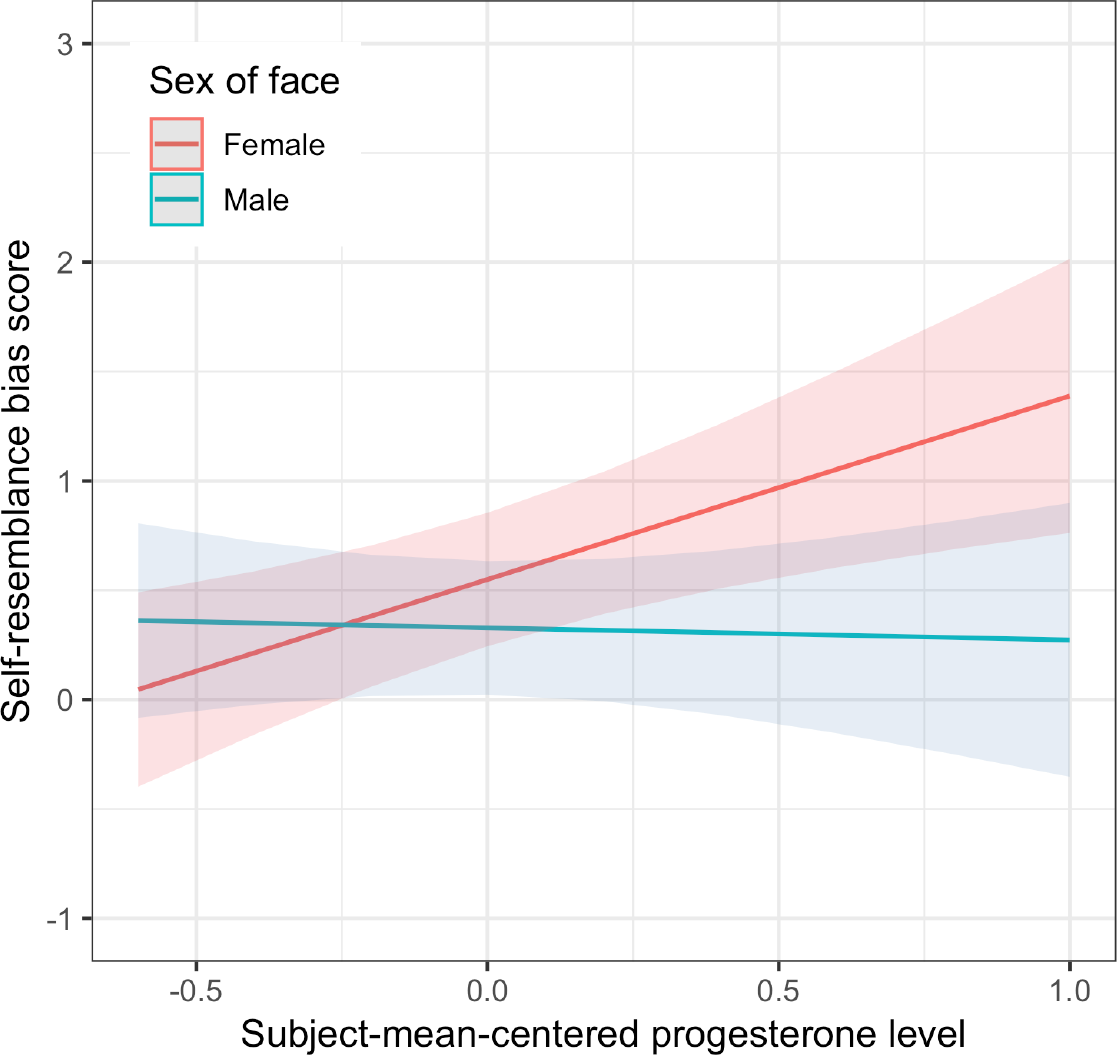
Model 2 showed a significant interaction between progesterone and face sex. The positive effect of progesterone on self-resemblance scores was greater for female than male faces.

## Study 2

Study 2 investigated whether women’s reported contact with and frequency of thought about male kin increased when conception risk was high in a sample of women not using hormonal contraceptives.

## Methods

### Participants

Two hundred and sixty-five women who were not using hormonal contraceptives (mean age=24.5 years, SD=5.1 years) and 106 women who were using hormonal contraceptives (mean age=24.5 years, SD=5.0 years) participated as part of a larger preregistered online diary study on conception risk and women’s mate preferences (https://osf.io/d3avf/). Inclusion criteria for taking part in this study were being single (i.e., not currently in a romantic relationship), heterosexual, and premenopausal, having a regular menstrual cycle (an average cycle length of at least 20 days), being younger than 50 years of age, and neither breastfeeding nor being pregnant now or in the three months prior to participation. Women who could not be clearly classed as hormonal contraceptive users or non-users (e.g., if they changed their contraceptive method during the course of the study or when they went off hormonal contraception less than three months prior to the study) or who lived with their parents were also excluded. Women who lived with their parents were excluded because their living arrangements will have directly influenced contact with male kin.

### Procedure

Initial questionnaires that were administered at the start of the project assessed factors such as relationship status and hormonal contraceptive use. Daily questionnaires were then administered in the evening after email and text message reminders. After 70 days, the daily questionnaires ended. In the daily questionnaires, women reported specific names or identifiers of people with whom they had had social contact (“With these people I had longer social contact (longer than 1 hour).”) and had thought about (“I thought about these people a lot and would have liked to see them.”). Every day, women could indicate whether their responses for a test session were dishonest or random, and such responses were discarded (less than 0.4% of days). In every third test session, women were asked to report the number of days since the onset of last period of menstrual bleeding (if it did not occur within the last five days of the diary, the next menstrual onset was assessed in a follow-up questionnaire). After finishing the daily diary, women completed a social network questionnaire that assessed their relationship to these individuals. Questionnaires were administered using the survey framework formr.org (Arslan, Walther & Tata, 2018). The full procedure (including variables not analysed here, is included in the supplemental materials).

### Analyses and results

The probability of being in the fertile window of the menstrual cycle was estimated for each test session using the backwards counting method following Gangestad et al. (2016) and Arslan et al. (2018). Following Arslan et al. (2018), for test sessions where we did not know the next menstrual onset, we counted backward from the last menstrual onset plus the reported average cycle length to estimate women’s probability of being fertile. In total, data for 12,931 test sessions from women not using hormonal contraceptives and 3,726 test sessions from women using hormonal contraceptives were available for analysis. Full results and analysis code for all analyses, as well as additional robustness checks, are given in our supplemental materials (see https://osf.io/f2hct/). None of these analyses showed credible evidence for a negative effect of conception risk on contact with and/or frequency of thoughts about male or other kin.

Our main outcome measure was whether or not women had contact with a biologically related man for more than an hour that day or thought about him a lot and would have liked to meet him. Lieberman et al.’s (2011) results predict that fertility will have a negative effect on contact with and thoughts about male kin in women not using hormonal contraceptives, but not in women using hormonal contraceptives. We tested this hypothesis by fitting a Bayesian probit regression model implemented in Stan (Carpenter et al., 2015) via the brms package (Bürkner, 2016). We had preregistered multilevel models with varying intercepts and varying slopes for the fertile window probability at the participant level (https://osf.io/73jha/). We adjusted for current menstruation and premenstrual phase (six days preceding menstrual onset). Hormonal contraception was included as a potential moderator. Our preregistered brms formula syntax in Wilkinson notation (Wilkinson & Rogers, 1973) is as follows:

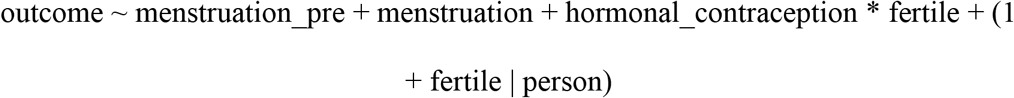

Results of this analysis are shown in Table 3. There was evidence of credible increases in contact with and thoughts about male kin during menstruation and the premenstrual phase. However, there was no credible evidence for a nonzero average effect of fertility in naturally cycling women. Women using hormonal contraceptives did not have more midcycle contact with male kin than women not using hormonal contraceptives, but rather had less, providing further evidence that contact with and thoughts about male kin do not decrease when conception risk is higher.

**Table 3.** Model summary of the main model. HDI = highest density interval, SD = standard deviation, cor = correlation

| Predictors | Whether person is a biologically related man |  |
| --- | --- | --- |
|  | *b* | *HDI (95%)* |
| Intercept | -3.11 | -3.53;-2.68 |
| Premenstrual phase | 0.16 | 0.06;0.27 |
| Menstruation | 0.13 | 0.01;0.25 |
| Hormonal contraception | -0.11 | -0.71;0.50 |
| Fertile phase | -0.33 | -1.39;0.60 |
| Hormonal contraception x fertile phase | -0.68 | -1.61;0.18 |
| Group-varying effects |  |  |
| SD(Woman Intercept) | 1.60 | 1.29;2.03 |
| SD(Woman Fertile phase) | 1.02 | 0.56;1.71 |
| cor(Intercept, Fertile Phase) | 0.31 | -0.44;0.81 |
| SD(Woman:Diary day Intercept) | 0.05 | 0.00;0.14 |
| Observations | 18652 coded interaction partners (1118 related men) on 8930 days across 240 women |  |

We also examined the varying slopes of the fertile phase on women’s contact with related men (see Figure 4.). For many women, no individual effects were estimable because of a lack of variation in the outcome. For four women (14.84%), the effect of the fertile phase on contact with related men was estimated to be positive and 95% credible intervals excluded zero (Figure 4).

**Figure 4.**
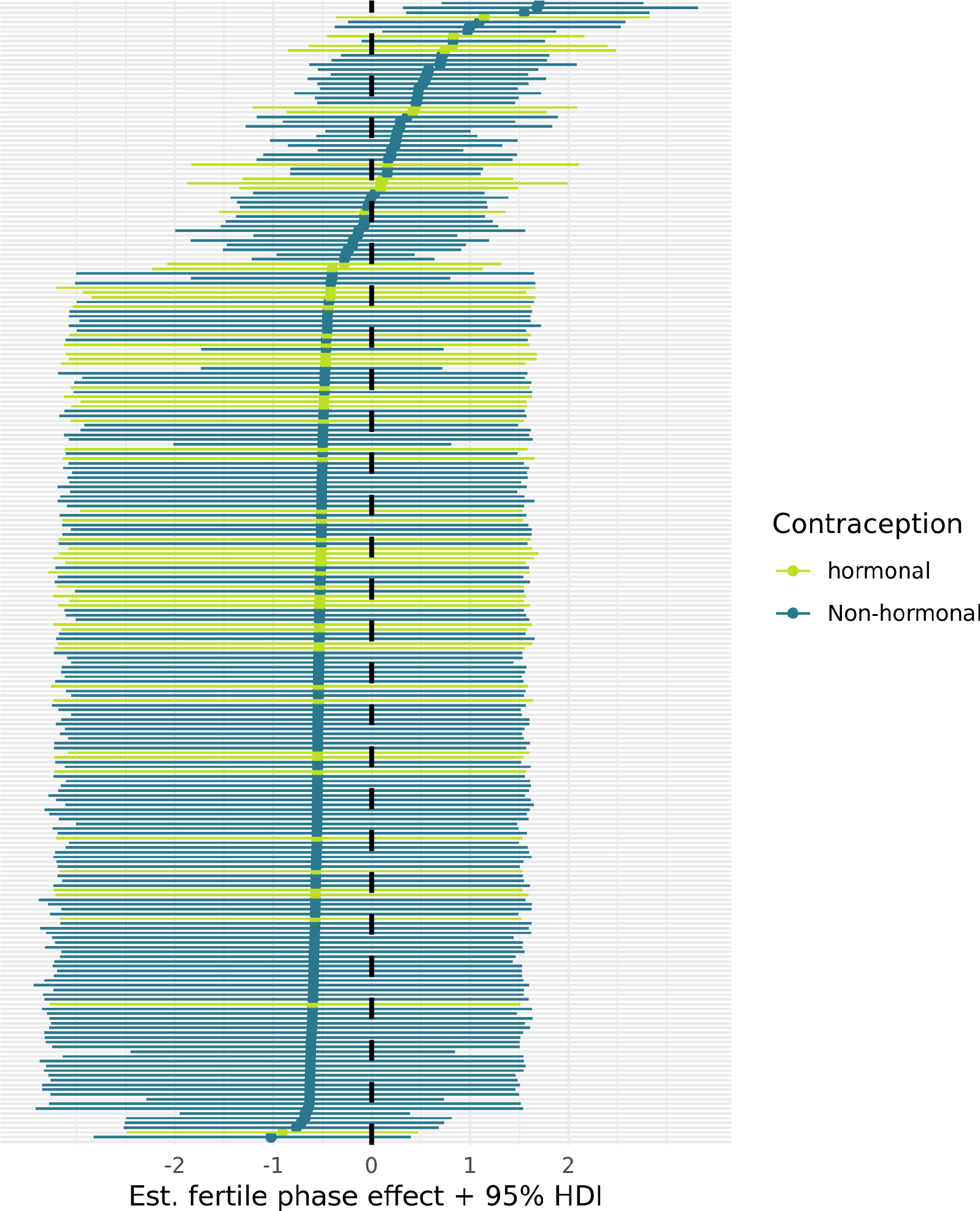
Forest plot of varying slopes of the fertile phase effect on women’s contact with related men. Each line and dot represent the estimate and 95% highest density interval (HDI) for the fertile phase effect on contact with related men for one woman, ordered by strength of the fertile phase change from top to bottom. Only where the horizontal line does not intersect with the vertical dashed line is there credible evidence for an intra-individual change. The women with the widest possible intervals lacked outcome variability, so their estimate simply reflects the mean slope with added uncertainty.

Next, we analysed contact with related men and thoughts about related men separately. The main effect of increased contact with male kin during women’s premenstrual phase (*b*=.21, 95% HDI [.07, .35]) and menstruation (*b*=.18, 95% HDI [.03, .34]) remained robust when examining actual social contact but disappeared when investigating thoughts about male kin.

## Discussion

In Study 1, we tested for evidence of hormonally regulated inbreeding avoidance in a longitudinal study of women’s responses to faces possessing kinship cues (i.e., self-resembling faces). By contrast with our predictions, we found no evidence that self-resemblance bias (i.e., the tendency to perceive self-resembling faces to be more attractive or trustworthy) decreased when fertility was high. In fact, in our analyses, self-resemblance bias when assessing men’s faces was *greater* when estradiol-to-progesterone ratio was higher (see Model 1). Since estradiol-to-progesterone ratio is positively correlated with conception risk during the menstrual cycle (Gangestad & Haselton, 2015; Puts et al., 2013), the observed positive effect of estradiol-to-progesterone ratio on self-resemblance bias when judging male faces is then in the opposite direction to what would be expected if inbreeding avoidance increased when fertility is high. Thus, our data from Study 1 do not support Lieberman et al.’s (2011) suggestion that inbreeding avoidance increases with conception risk during the menstrual cycle.

Although we found no evidence that self-resemblance bias was weaker when fertility was high, women did (on average) judge self-resembling faces to be more trustworthy and attractive than non-self-resembling faces. This tendency to perceive self-resembling faces more positively than would be expected by chance alone replicates results from previous research (DeBruine, 2002, 2004, 2005). We also found that women’s self-resemblance bias tended to be greater when judging women’s faces than when judging men’s faces, although this effect of sex of face was only significant in one analysis^1^. Nonetheless, in both analyses, the direction of the effect of face sex is consistent with both previous research (DeBruine, 2004) and the proposal that inbreeding avoidance acts as a brake on the self-resemblance bias in social judgments of faces (DeBruine et al., 2008).

In a cross-sectional study, DeBruine et al. (2005) reported that self-resemblance bias increased when progesterone levels were relatively high when assessing women’s, but not men’s, faces. Evidence for such an effect in our sample was mixed. Both of our analyses found that the self-resemblance bias was stronger when progesterone was higher. Results from Model 2, but not Model 1, suggested that this positive effect of progesterone on self-resemblance bias scores was driven by responses to female faces. DeBruine et al. (2005) suggested that stronger self-resemblance bias for women’s faces when progesterone is high could function to increase bonding with female kin when raised progesterone prepares the body for pregnancy and support from kin may be particularly beneficial. However, in the current study, we also found that self-resemblance bias was weaker when estradiol was higher. Both estradiol and progesterone are elevated during pregnancy (Johnson, 2007). Thus, that progesterone and estradiol have opposite effects on self-resemblance bias does not straightforwardly support DeBruine et al.’s (2005) proposal that stronger self-resemblance bias when progesterone is high reflects hormonal regulation of responses to kinship cues that evolved to increase bonding with kin during pregnancy.

In Study 2, we investigated whether women’s reported contact with and frequency of thoughts about male kin tracked changes in their conception risk over the menstrual cycle. We found no evidence that contact with male kin or frequency of thoughts about male kin decreased when conception risk was high in women not using hormonal contraceptives. In fact, our quasi-control group of women using hormonal contraceptives showed a greater mid-cycle increase in aversion to male kin compared to women not using hormonal contraceptives.

In summary, the main finding from Study 1 was that self-resemblance bias scores for male faces were positively (rather than negatively) related to estradiol-to-progesterone ratio (a well-validated proxy measure of conception risk). This pattern of results directly contradicts (i.e., is in the opposite direction to what would be predicted by) Lieberman et al.’s (2011) hypothesis that inbreeding-avoidance behaviors increase before ovulation. The main finding from Study 2 was that women using hormonal contraceptives showed mid-cycle increase in aversion to male kin, whereas women not using hormonal contraceptives did not. Again, this pattern of results directly contradicts Lieberman et al.’s (2011) hypothesis that inbreeding-avoidance behaviors increase during ovulation. Thus, despite both being larger-scale studies than the original Lieberman et al. (2011) study, neither of our studies found any evidence that inbreeding-avoidance behaviors increase when women are fertile. Our results then raise the possibility that the significant finding reported in the original Lieberman et al. (2011) study is not robust.

## Supplemental Material

### Participants

The recruitment took place between June 2016 and January 2017 through various online channels (e.g. online platform psytests.de, advertisement on okCupid.com and Facebook and mass mailing lists of university students) as well as direct invitations of suitable candidates taking part in previous lab studies. Data collection ended in May 2017.

The incentives for taking part in the study were either direct payment of participants with an amount ranging from 25€ up to 45€^2^. Alternatively, participants had the chance of winning prizes with a total value of 2,000€^3^. Students of the University of Goettingen were also able to earn course credit. For all three rewards, the amount of credit, money, or lots depended on the regularity of participation. At the end of the study, every participant received a personalised graphical feedback as a further incentive.

The final sample consists of 371 women with 265 naturally cycling women and 106 women taking hormonal contraceptives. Age ranges from 18 to 43 years (M = 24.48, SD = 5.08) for naturally cycling women and from 18 to 42 years (M = 24.48, SD = 5.03) for hormonal contraceptive users. Table 1 lists a detailed description of participants. There were no group differences in demographic variables except that women taking hormonal contraceptives had their first sexual intercourse earlier than naturally cycling women on average.

**Table S1.** Descriptive of naturally cycling women and women taking hormonal contraceptives.

|  | Naturally cycling women |  |  |  | Hormonal contraceptive user |  |  |  | p |
| --- | --- | --- | --- | --- | --- | --- | --- | --- | --- |
|  | M | SD | Min | Max | M | SD | Min | Max |  |
| Age | 24.48 | 5.08 | 18 | 43 | 24.58 | 5.03 | 18 | 42 | NS |
| Cycle length | 28.67 | 3.13 | 20 | 40 | 28.08 | 3.39 | 21 | 49 | NS |
| Education | 14.49 | 5.14 | 0 | 26 | 14.55 | 4.38 | 0 | 26 | NS |
| Age at first sex | 17.58 | 2.79 | 13 | 29 | 16.79 | 2.12 | 13 | 26 | .006 |
| Number of sexual partners | 8.27 | 12.88 | 0 | 100 | 8.19 | 9.90 | 0 | 70 | NS |

*Note.* NS = not significant.**Study structure**

### Study structure

Women participated in an online study named “Alltag und Sexualität [Daily Life and Sexuality]” implemented using the survey framework formr.org (Arslan, Walther & Tata, 2018). The study was introduced as an online diary which aimed to examine the interaction of sexuality, psychological well-being, experience of romantic relationships with everyday experiences. The study had six main stages. Figure 1 facilitates understanding of the study’s structure.

The first step was to fill out a demographic questionnaire which served as an initial screening phase for suitable participants. Afterwards a personality questionnaire irrelevant to the current study followed.

A day after these surveys, women started the online diary. Over a period of 70 days, women received an online invitation via email at 5 pm. This online diary could be filled out until 3 am the following day and included questions about their everyday life. Items were randomised within grouped blocks. Important items were shown every day while items of lower importance randomly appeared 20-70% of the time.

The fourth step was a social network questionnaire only sent to the single women. In the diary, participants had the chance to list persons that they met or thought about in form of acronyms or nicknames. For up to ten of these names – if they had been mentioned three or more times – further questions were asked starting with the least frequently mentioned person in ascending order. If the focal person remembered who she meant by the name, we requested to know the gender and relationship towards the anchor. In the case of unrelated men, further questions followed including the perceived attractiveness as a short- and long-term mate and several other characteristics.

Finally, a follow-up questionnaire assessed whether important changes during the diary occurred. Additionally, the next menstrual onset was assessed for women reporting their menstruation more than five days before the end of the diary.

**Figure S1.**
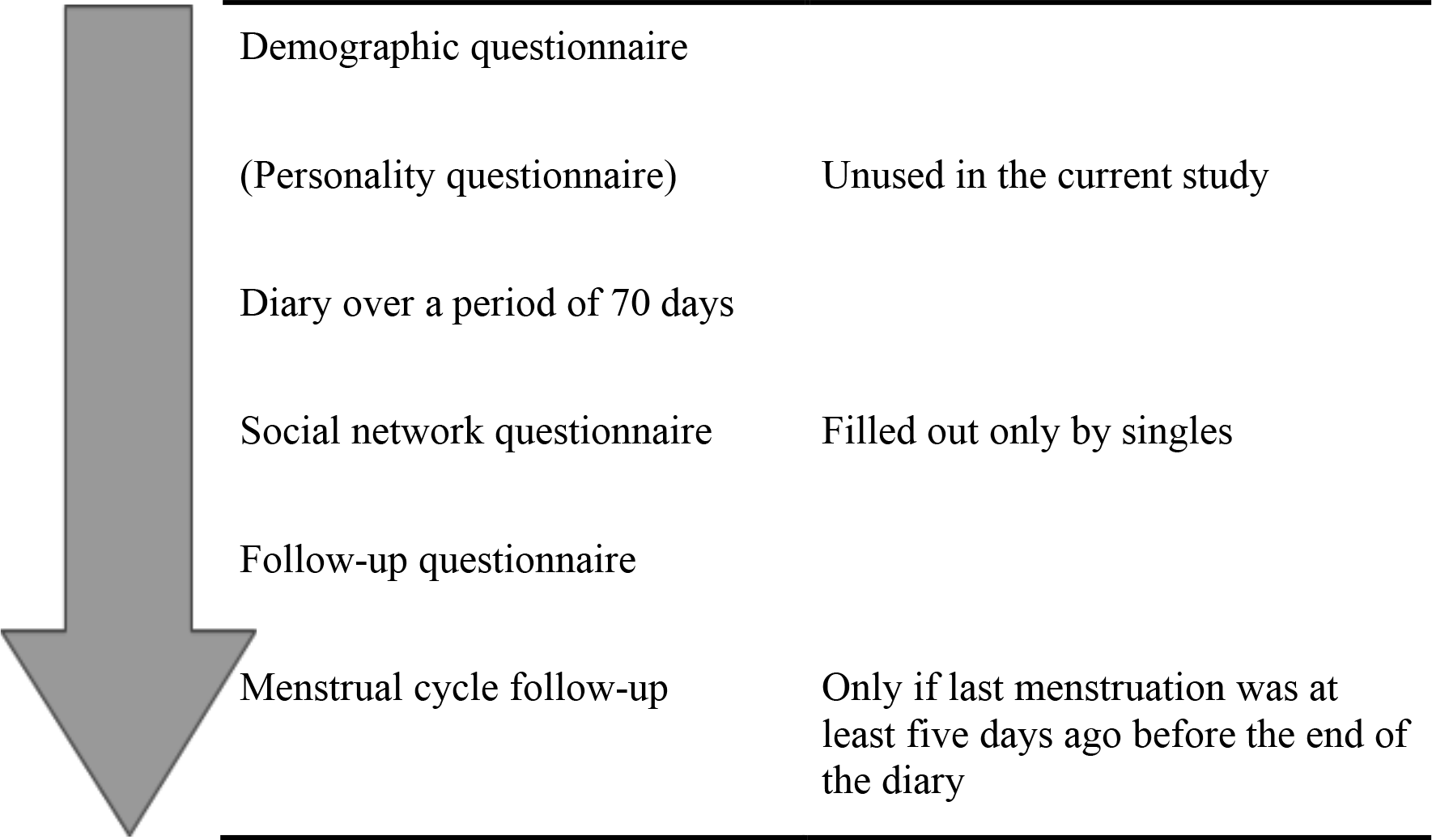
Procedure.

### Footnotes

1 In Model 1, the estimate for the effect of face sex was −0.21 (*p*=.068). The corresponding effect in Model 2 was −0.22 (*p*=.003).

2 Only women fulfilling certain sample criteria are offered direct payment. Those were being under the age of 50, having a heterosexual orientation, a regular menstruation and being pre-menopausal as well as having not taken any hormonal or psychoactive medication and no hormonal contraception in the last three months. Additionally, women were only paid if they were not trying to get pregnant or were pregnant and/or breast feeding within the last three months.

3 The prizes of the lottery included an iPhone, an iPad and forty 20€ Amazon coupons.

## References

Arslan, R. C., Schilling, K. M., Gerlach, T. M., & Penke, L. (2018). Using 26,000 diary entries to show ovulatory changes in sexual desire and behavior. Journal of Personality and Social Psychology, Advance Online Publication. https://doi.org/10.1037/pspp0000208

Arslan, R. C., Walther, M., & Tata, C. (2018). formr: A study framework allowing for automated feedback generation and complex longitudinal experience sampling studies using R. PsyArXiv. https://doi.org/10.31234/osf.io/pjasu

Bao, A. M., Liu, R. Y., van Someren, E. J., Hofman, M. A., Cao, Y. X., & Zhou, J. N. (2003). Diurnal rhythm of free estradiol during the menstrual cycle. European Journal of Endocrinology, 148(2), 227–232. Retrieved from https://eje.bioscientifica.com/view/journals/eje/148/2/227.xml

Barr, D. J. (2013). Random effects structure for testing interactions in linear mixed-effects models. Frontiers in Psychology, 4(328). https://doi.org/10.3389/fpsyg.2013.00328

Barr, D. J., Levy, R., Scheepers, C., & Tily, H. J. (2013). Random effects structure for confirmatory hypothesis testing: Keep it maximal. Journal of Memory and Language, 68(3), 255–278. https://doi.org/10.1016/j.jml.2012.11.001

Bates, D., Maechler, M., Bolker, B., & Walker, S. (2015). Fitting Linear Mixed-Effects Models Using lme4. Journal of Statistical Software, 67(1), 1–48.https://doi.org/10.18637/jss.v067.i01

Blake, K. R., Dixson, B. J. W., O'Dean, S. M., & Denson, T. F. (2016). Standardized protocols for characterizing women's fertility: A data-driven approach. Hormones and Behavior, 81, 74–83. https://doi.org/10.1016/j.yhbeh.2016.03.004

Bürkner, P.-C. (2017). brms: An R Package for Bayesian Multilevel Models Using Stan. Journal of Statistical Software, 80(1), 1–28. https://doi.org/10.18637/jss.v080.i01

Carpenter, B., Gelman, A., Hoffman, M. D., Lee, D., Goodrich, B., Betancourt, M., Brubaker, M., Guo, J. Li, P. & Riddell, A. (2017). Stan: A Probabilistic Programming Language. Journal of Statistical Software, 76(1), 1–32. https://doi.org/10.18637/jss.v076.i01

DeBruine, L. M. (2002). Facial resemblance enhances trust. Proceedings of the Royal Society B: Biological Sciences, 269, 1307–1312. https://doi.org/10.1098/rspb.2002.2034

DeBruine, L. M. (2004). Facial resemblance increases the attractiveness of same-sex faces more than other-sex faces. Proceedings of the Royal Society B: Biological Sciences, 271, 2085–2090. https://doi.org/10.1098/rspb.2004.2824

DeBruine, L. M. (2005). Trustworthy but not lust-worthy: context-specific effects of facial resemblance. Proceedings of the Royal Society B: Biological Sciences, 272, 919–922. https://doi.org/10.1098/rspb.2004.3003

DeBruine, LM (2018). DeBruine/Webmorph: Beta release 2 (Version v0.0.0.9001). Zenodo. https://doi.org/10.5281/zenodo.1162670

DeBruine, L. M., Jones, B. C., & Perrett, D. I. (2005). Women's attractiveness judgments of self-resembling faces change across the menstrual cycle. Hormones and Behavior, 47, 379–383. https://doi.org/10.1016/j.yhbeh.2004.11.006

DeBruine, L. M., Jones, B. C., Little, A. C., & Perrett, D. I. (2008). Social perception of facial resemblance in humans. Archives of Sexual Behavior, 37, 64–77. https://doi.org/10.1007/s10508-007-9266-0

Gangestad, S. W., Garver-Apgar, C. E., Simpson, J. A., & Cousins, A. J. (2007). Changes in women's mate preferences across the ovulatory cycle. Journal of Personality and Social Psychology, 92, 151–163. https://doi.org/10.1037/0022-3514.92.1.151

Gangestad, S. W., & Haselton, M. G. (2015). Human estrus: implications for relationship science. Current Opinion in Psychology, 1, 45–51. https://doi.org/10.1016/j.copsyc.2014.12.007

Gangestad, S. W., Simpson, J. A., Cousins, A. J., Garver-Apgar, C. E., & Christensen, P. N. (2004). Women's preferences for male behavioral displays change across the menstrual cycle. Psychological Science, 15, 203–207. https://doi.org/10.1111/j.0956-7976.2004.01503010.x

Gangestad, S. W., & Thornhill, R. (2008). Human oestrus. Proceedings of the Royal Society B: Biological Sciences, 275, 991–1000. https://doi.org/10.1098/rspb.2007.1425

Gangestad, S. W., Haselton, M. G., Welling, L. L. M., Gildersleeve, K., Pillsworth, E. G., Burriss, R. P., … Puts, D. A. (2016). How valid are assessments of conception probability in ovulatory cycle research? Evaluations, recommendations, and theoretical implications. Evolution and Human Behavior, 37(2), 85–96. https://doi.org/10.1016/j.evolhumbehav.2015.09.001

Gildersleeve, K., Haselton, M. G., & Fales, M. R. (2014). Do women's mate preferences change across the ovulatory cycle? A meta-analytic review. Psychological Bulletin, 140, 1205–1259. https://doi.org/10.1037/a0035438

Johnson MH (2007) Essential Reproduction (Blackwell Publishing) 6 Ed.

Jones, B. C., DeBruine, L. M., Perrett, D. I., Little, A. C., Feinberg, D. R., & Law Smith, M. J. (2008). Effects of menstrual cycle phase on face preferences. Archives of Sexual Behavior, 37, 78–84. https://doi.org/10.1007/s10508-007-9268-y

Jones, B. C., Hahn, A. C., & DeBruine, L. M. (in press). Ovulation, sex hormones, and women’s mating psychology. Trends in Cognitive Sciences.

Jones, B. C., Hahn, A. C., Fisher, C. I., Wang, H., Kandrik, M., Han, C., … DeBruine, L. M. (2018a). No Compelling Evidence that Preferences for Facial Masculinity Track Changes in Women’s Hormonal Status. Psychological Science, 29(6), 996–1005. https://doi.org/10.1177/0956797618760197

Jones, B. C., Hahn, A. C., Fisher, C. I., Wang, H., Kandrik, M., Lee, A. J., … DeBruine, L. M. (2018b). Hormonal correlates of pathogen disgust: testing the compensatory prophylaxis hypothesis. Evolution and Human Behavior, 39(2), 166–169. https://doi.org/10.1016/j.evolhumbehav.2017.12.004

Jones, B. C., Hahn, A. C., Fisher, C. I., Wang, H., Kandrik, M., & DeBruine, L. M. (2018c). General sexual desire, but not desire for uncommitted sexual relationships, tracks changes in women’s hormonal status. Psychoneuroendocrinology, 88, 153–157. https://doi.org/10.1016/j.psyneuen.2017.12.015

Jünger, J., Gerlach, T. M., & Penke, L. (2018a). No evidence for ovulatory cycle shifts in women’s preferences for men’s behaviors in a pre-registered study. https://doi.org/10.31234/osf.io/7g3xc

Jünger, J., Motta-Mena, N. V., Cardenas, R., Bailey, D., Rosenfield, K. A., Schild, C., … Puts, D. A. (2018b). Do women's preferences for masculine voices shift across the ovulatory cycle? Hormones and Behavior, 106, 122–134. https://doi.org/10.1016/j.yhbeh.2018.10.008

Kuznetsova, A., Brockhoff, P. B., & Christensen, R. H. B. (2017). lmerTest Package: Tests in Linear Mixed Effects Models. Journal of Statistical Software, 82(13), 1–26. https://doi.org/10.18637/jss.v082.i13

Lieberman, D., Tooby, J., & Cosmides, L. (2007). The architecture of human kin detection. Nature, 445(7129), 727–731. https://doi.org/10.1038/nature05510

Lieberman, D., Pillsworth, E. G., & Haselton, M. G. (2011). Kin affiliation across the ovulatory cycle. Psychological Science, 22(1), 13–18. https://doi.org/10.1177/0956797610390385

Marcinkowska, U. M., Galbarczyk, A., & Jasienska, G. (2018). La donna è mobile? Lack of cyclical shifts in facial symmetry, and facial and body masculinity preferences—A hormone based study. Psychoneuroendocrinology, 88, 47–53. https://doi.org/10.1016/j.psyneuen.2017.11.007

Papacosta, E., & Nassis, G. P. (2011). Saliva as a tool for monitoring steroid, peptide and immune markers in sport and exercise science. Journal of Science and Medicine in Sport, 14(5), 424–434. http://dx.doi.org/10.1016/j.jsams.2011.03.004

Penton-Voak, I. S., & Perrett, D. I. (2000). Female preference for male faces changes cyclically: Further evidence. Evolution and Human Behavior, 21, 39–48. https://doi.org/10.1016/S1090-5138(99)00033-1

Penton-Voak, I. S., Perrett, D. I., Castles, D. L., Kobayashi, T., Burt, D. M., Murray, L. K., & Minamisawa, R. (1999). Menstrual cycle alters face preference. Nature, 399, 741. https://doi.org/10.1038/21557

Puts, D. A., Bailey, D. H., Cárdenas, R. A., Burriss, R. P., Welling, L. L. M., Wheatley, J. R., & Dawood, K. (2013). Women's attractiveness changes with estradiol and progesterone across the ovulatory cycle. Hormones and Behavior, 63(1), 13–19. https://doi.org/10.1016/j.yhbeh.2012.11.007

R Core Team. (2018). R: A language and environment for statistical computing (Version 3.5.0). Vienna, Austria: R Foundation for Statistical Computing. Retrieved from https://www.R-project.org/

Tiddeman, B. P., Burt, D. M., & Perrett, D. I. (2001). Prototyping and transforming facial textures for perception research. IEEE Computer Graphics and Applications, 21, 42–50. https://doi.org/10.1109/38.946630

Veldhuis, J. D., Christiansen, E., Evans, W. S., Kolp, L. A., Rogol, A. D., & Johnson, L. (1988). Physiological Profiles of Episodic Progesterone Release During the Midluteal Phase of the Human Menstrual Cycle: Analysis of Circadian and Ultradian Rhythms, Discrete Pulse Properties, and Correlations With Simultaneous Luteinizing Hormone Release. The Journal of Clinical Endocrinology & Metabolism, 66(2), 414–421. https://doi.org/10.1210/jcem-66-2-414

Wilkinson, G. N., & Rogers, C. E. (1973). Symbolic Description of Factorial Models for Analysis of Variance. Journal of the Royal Statistical Society. Series C (Applied Statistics), 22(3), 392–399. https://doig.org/10.2307/2346786

Wood, W., Kressel, L., Joshi, P. D., & Louie, B. (2014). Meta-analysis of menstrual cycle effects on women's mate preferences. Emotion Review, 6, 229–249. https://doi.org/10.1177/1754073914523073

## References

Arslan, R. C., Walther, M., & Tata, C. (2018). formr: A study framework allowing for automated feedback generation and complex longitudinal experience sampling studies using R. PsyArXiv. doi:10.31234/osf.io/pjasu

